# A multiplexed RT-PCR Assay for Nanopore Whole Genome Sequencing of Tilapia lake virus (TiLV)

**DOI:** 10.1101/2023.04.24.537954

**Authors:** Jerome Delamare-Deboutteville, Watcharachai Meemetta, Khaettareeya Pimsannil, Pattiya Sangpo, Han Ming Gan, Chadag Vishnumurthy Mohan, Ha Thanh Dong, Saengchan Senapin

**Author notes:** Corresponding authors: J. Delamare-Deboutteville, S. Senapin. these authors contributed equally to this work (Jerome Delamare-Deboutteville, Watcharachai Meemetta).

## Abstract

Tilapia lake virus (TiLV) is a highly contagious viral pathogen that affects tilapia, a globally significant and affordable source of fish protein. To prevent the introduction and spread of TiLV and its impact, there is an urgent need for increased surveillance, improved biosecurity measures, and continuous development of effective diagnostic and rapid sequencing methods. In this study, we have developed a multiplexed RT-PCR assay that can amplify all ten complete genomic segments of TiLV from various sources of isolation. The amplicons generated using this approach were immediately subjected to real-time sequencing on the Nanopore system. By using this approach, we have recovered and assembled 10 TiLV genomes from total RNA extracted from naturally TiLV-infected tilapia fish, concentrated tilapia rearing water, and cell culture. Our phylogenetic analysis, consisting of more than 36 TiLV genomes from both newly sequenced and publicly available TiLV genomes, provides new insights into the high genetic diversity of TiLV. This work is an essential steppingstone towards integrating rapid and real-time Nanopore-based amplicon sequencing into routine genomic surveillance of TiLV, as well as future vaccine development.

## Introduction

Tilapia lake virus disease (TiLVD) is a highly contagious viral disease that affects tilapia, an important and affordable fish protein source produced in aquaculture globally. TiLV has rapidly spread since, now reported in over 18 countries ^1–4^, posing a significant threat to tilapia production and the livelihoods of farmers who rely on tilapia farming for income and food security ^5^. To mitigate the introduction and spread of TiLV and its impacts, it is imperative to implement surveillance, improved biosecurity measures, farming practices and continuous development of effective diagnostic and rapid sequencing methods.

The TiLV genome consists of 10 segments that complicate its genome sequencing process thus precluding high mass-scale genome sequencing efforts to be undertaken. The first TiLV genome was sequenced using a shotgun transcriptome approach on an Illumina sequencing platform ^6^. The genomes of TiLV were also sequenced using the Sanger sequencing technique ^7,8^. Recently, a similar approach using shotgun metagenomics was used to generate the near complete genome of a TiLV isolate causing mass-mortality in tilapia farmed in Bangladesh ^9^. Shotgun metagenomics involved the random sequencing of all RNA fragments, i.e., TiLV-positive tilapia liver samples without any enrichment of mRNA; followed by bioinformatics analysis to identify and assemble the 10-segments of TiLV genome present in the sample. This approach is not scalable as the RNA library preparation cost is higher and most of the sequencing data will belong to the host, requiring high sequencing depth to successfully assemble the TiLV-derived contigs. To address these challenges and improve sequencing effectiveness, some approaches have been explored. One such approach involves the propagation of viruses in cell culture prior to sequencing ^10^. Additionally, enrichment through single RT-PCR amplification of the TiLV 10 segments before subjecting them to Illumina sequencing has also been employed ^3^. Nevertheless, the use of Illumina technology necessitates a significant investment in infrastructure, which hinders rapid on-site deployment and real-time sequencing.

In recent years, Oxford Nanopore Technologies (ONT) have become more commonly used to sequence part(s) or whole genomes of pathogens affecting aquatic animals ^11^. Nanopore sequencing offers several advantages, including high throughput, portability, cost-effectiveness, and real-time sequencing, which can greatly facilitate the detection and sequencing of viral genomes in remote locations ^12^. Rapid amplicon-based reverse transcription polymerase chain reaction (RT-PCR) assays coupled with Nanopore technologies can provide a sensitive and specific means of detecting and genotyping TiLV in field samples, allowing for fine epidemiological surveillance and timely management and control of outbreaks ^13^. To date, Nanopore has been used to sequence the genomes of non-segmented fish viruses such as infectious spleen and kidney necrosis virus (ISKNV) ^12^, salmonid alphavirus (SAV) and infectious salmon anaemia virus (ISAV) ^11^.

In this study, for the first time, we designed a new method for TiLV whole genome sequencing using singleplex and multiplex amplicon-based RT-PCR protocols coupled with Minion Nanopore sequencing. These novel tools enable real-time diagnosis and characterization of TiLV genomes, thereby facilitating improved surveillance and effective control measures in tilapia aquaculture.

## Methods

### Ethics declarations

The authors confirm that the ethical policies of the journal, as noted on the journal’s submission guidelines page, have been adhered to. No ethical approval was required as no animals were used in this study. Virus sequences were generated from archived samples.

### Primer design

TiLV primers targeting 10 complete genome segments (containing conserved sequences at 5’ and 3’ termini) were designed based on sequences of the Israel strain Til-4-2011 (GenBank accession no. KU751814 to KU751823) ^6^ (Table 1).

**Table 1.**
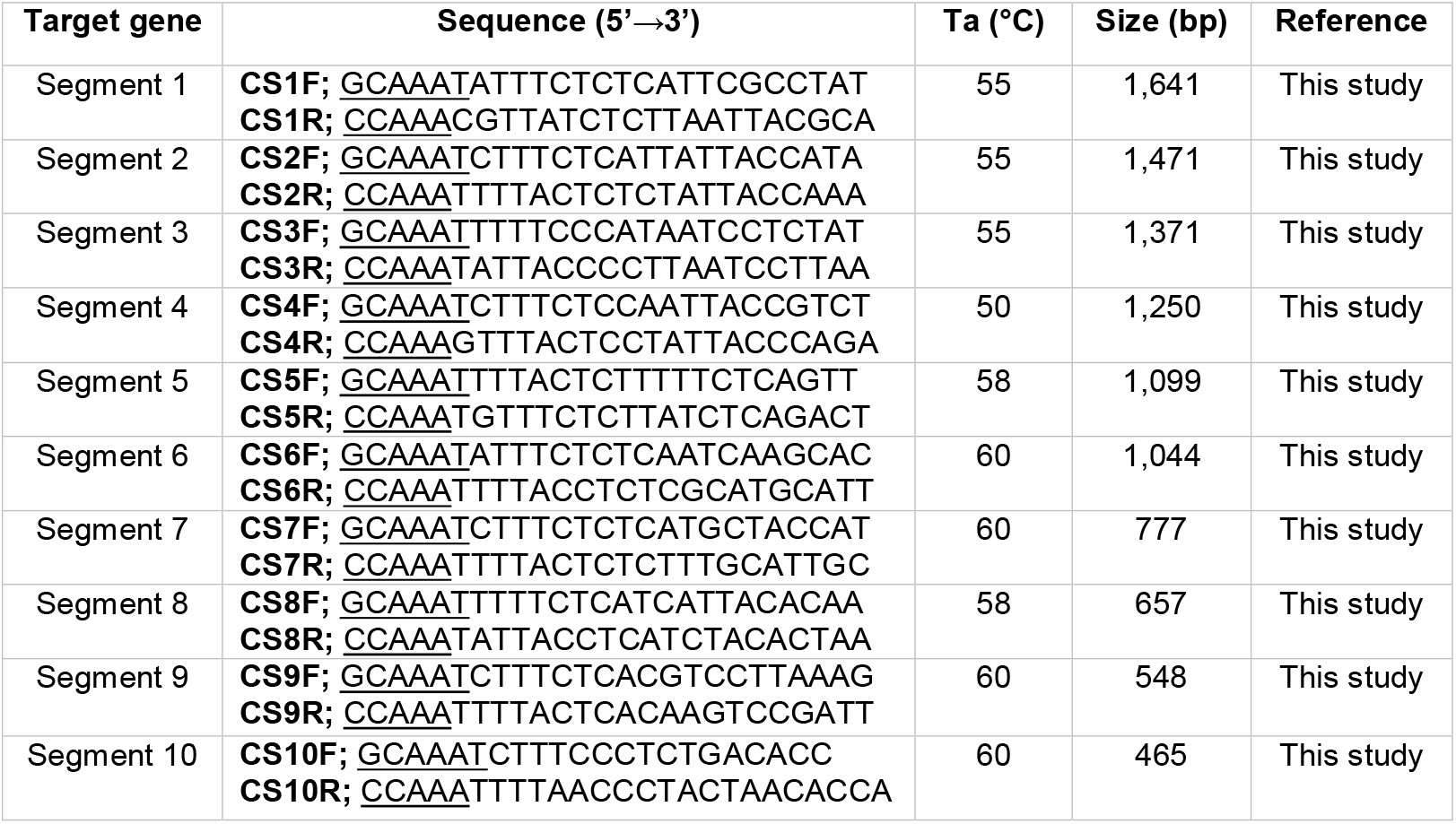
TiLV primers designed in this study. Underlines represent conserved TiLV genomic sequences at the 5’ and 3’ end of each segment.

### Samples, total RNA extraction, and TiLV quantification

RNA template (N = 10) for TiLV genome sequence amplification and analysis was prepared from tissues of TiLV infected Nile tilapia (*Oreochromis niloticus*), red tilapia (*Oreochromis* spp.), TiLV isolates propagated in E-11 cell culture, and concentrated water samples from fish ponds (Table 2). Extraction of total RNA used the Trizol reagent (Invitrogen) according to the manufacturer’s instruction followed by spectrophotometry-based quantification measuring the absorbance at OD_260 nm_ and OD_280 nm_. TiLV quantification by probe-based qPCR assays of the 10 samples were performed based on segment 9 ^14^ and segment 1 (this study) (Supplemental Table 1).

**Table 2.** Background information of TiLV strains reported in study as well as strains with publicly available genomes.

| # | Sequence name (strain) | Country of origin | Collection date | Fish host | Tissues | Accession number | Reference | Sequencing Technology |
| --- | --- | --- | --- | --- | --- | --- | --- | --- |
| 1 | B1-1_m | TH | 2021 | Red tilapia | L, K, S | SRR24222694 | This study | Nanopore |
| 2 | B1-2_m | TH | 2021 | Red tilapia | L, K, S | SRR24222693 | This study | Nanopore |
| 3 | FM2_m | TH | 2019 | Nile tilapia | L, K, S | SRR24222689 | This study | Nanopore |
| 4 | Nk_m | TH | 2019 | Nile tilapia | L, B | SRR24222688 | This study | Nanopore |
| 5 | A1-2_m | TH | 2021 | Red tilapia | L, K, S | SRR24222687 | This study | Nanopore |
| 6 | A1-3_s | TH | 2021 | Red tilapia | L, K, S | SRR24222682 | This study | Nanopore |
| 7 | A1-3_m | TH | 2021 | Red tilapia | L, K, S | SRR24222686 | This study | Nanopore |
| 8 | D1-2_s | TH | 2021 | Red tilapia | L, K, S | SRR24222692 | This study | Nanopore |
| 9 | D1-2_m | TH | 2021 | Red tilapia | L, K, S | SRR24222685 | This study | Nanopore |
| 10 | Ri_m | TH | 2020 | Nile tilapia | L | SRR24222684 | This study | Nanopore |
| 11 | Cell_line_s | TH | 2019 | Red tilapia | E-11 cells | SRR24222690 | This study | Nanopore |
| 12 | Cell_line_m | TH | 2019 | Red tilapia | E-11 cells | SRR24222683 | This study | Nanopore |
| 13 | RiverWater_s | TH | 2021 | Red tilapia (water) | na | SRR24222691 | This study | Nanopore |
| 14 | IL-2011 (Til-4-2011) | IL | 1 May 2011 | Hybrid tilapia | na | KU751814-823 | <sup>23</sup> | Illumina |
| 15 | TH-2013 | TH | Jan 2013 | Nile tilapia | Fe | MN687685-695 | <sup>24</sup> | Sanger |
| 16 | TH-2014 | TH | Aug 2014 | Nile tilapia | Fr | MN687695-704 | <sup>8</sup> | Sanger |
| 17 | TH-2015 | TH | Aug 2015 | Nile tilapia | Fi | MN687705-714 | <sup>8</sup> | Sanger |
| 18 | TH-2016-CU | TH | Dec 2016 | Nile tilapia | Ju | MN687715-724 | <sup>8</sup> | Sanger |
| 19 | TH-2016-CN | TH | Dec 2016 | Red tilapia hybrid | Fi | MN687725-734 | <sup>8</sup> | Sanger |
| 20 | TH-2016 (TV1) | TH | 2016 | Red tilapia | na | KX631921-930 | <sup>7</sup> | Sanger |
| 21 | TH-2017 | TH | 2017 | Nile tilapia | na | MN687735-744 | <sup>8</sup> | Sanger |
| 22 | TH-2018-N | TH | Jul 2018 | Red tilapia | Fi | MN687745-754 | <sup>8</sup> | Sanger |
| 23 | TH-2018-K | TH | Aug 2018 | Nile tilapia | Ju | MN687755-764 | <sup>8</sup> | Sanger |
| 24 | TH-2018 (WVL18053-01A) | USA | Apr 2018 | Nile tilapia |  | MH319378-387 | <sup>10</sup> | Illumina |
| 25 | TH-2019 | TH | Feb 2019 | Nile tilapia | Fi | MN687765-774 | <sup>8</sup> | Sanger |
| 26 | EC-2012 (EC-2012) | EC | 2012 | Nile tilapia | na | MK392372-381 | <sup>25</sup> | Illumina |
| 27 | PE-2018 (F3-4) | PE | 2018 | Nile tilapia | na | MK425010-019 | <sup>26</sup> | Sanger |
| 28 | US-2019 (WVL19031-01A) | US | 2018 | Nile tilapia | na | MN193513-522 | unpublished* | Illumina |
| 29 | US-2019 (WVL19054) | US | 2019 | Nile tilapia | na | MN193523-532 | unpublished* | Illumina |
| 30 | BD-2017 | BD | 2017 | Nile tilapia | na | MN939372-381 | <sup>9</sup> | Illumina |
| 31 | BD-2019-E1 | BD | 2019 | Nile tilapia | na | MT466447-456 | <sup>27</sup> | Sanger |
| 32 | BD-2019-E3 | BD | 2019 | Nile tilapia | na | MT466457-466 | <sup>27</sup> | Sanger |
| 33 | BD-2017-181 | BD | 2017 | Nile tilapia | na | MT466437-446 | <sup>27</sup> | Sanger |
| 34 | IND-2018 | IND | 2017 | Nile tilapia | na | MZ297923-932 | unpublished* | Illumina |
| 35 | VN (HB196-VN-2020) | VN | 2020 | Nile tilapia | na | ON376572-581 | <sup>3</sup> | Illumina; Sanger |
| 36 | VN (RIA2-VN-2019) | VN | 2019 | Nile tilapia | na | ON376582-591 | <sup>3</sup> | Illumina; Sanger |

Country of origin: TH, Thailand; IL, Israel; EC, Ecuador; PE, Peru; US, USA; BD, Bangladesh; IND, India; VN, Vietnam Fish host: Red tilapia, *Oreochromis* sp.; Nile tilapia, *Oreochromis niloticus*; Hybrid tilapia *Oreochromis niloticus* x *Oreochromis aureus*; L, Liver; K, Kidney; S, Spleen; B, Brain; Fe, fertilized eggs; Fr, fry; Fi, fingerlings; Ju, juveniles; na, info not available; \*, direct submission. Sequence name for samples #1-13: \_s (singleplex PCR), \_m (multiplex PCR).

## Development of singleplex one-step RT-PCR for the enrichment of TiLV genome

The efficiency of the designed TiLV primers and their optimal annealing temperatures (Ta) were investigated by one-step gradient RT-PCR assays with the range of Ta from 50 to 60°C. RT-PCR reaction mixture (25 μl) comprised of 0.5 μl of SuperScript III RT/Platinum Taq Mix (Invitrogen catalog no. 12574-018), 12.5 μl of 2X reaction mix, 2 μl of RNA template (100 ng/ μl), 1 μl of 10 μM each primer pair, and distilled water. The temperature profile included a reverse transcription (RT) step at 50°C for 30 min, heat-inactivation of RT enzyme at 94°C for 2 min, 30 cycles of denaturation at 94°C for 30 sec, annealing step for 30 sec, extension 72°C for 1 min-2 min (1min/kb), and a final extension at 72°C for 2 min. Amplified products of the singleplex RT-PCR (sPCR) were analyzed by agarose gel electrophoresis. Four RNA templates were used in this assay (Table 2).

## Development of multiplex RT-PCR to streamline the PCR enrichment of TiLV genome

Two multiplex PCR (mPCR) reactions were developed to reduce the number of PCR reactions from 10 to only two reactions per sample. The primers were divided into two sets based on their annealing temperatures similarity. Reaction 1 employs primers for segment 1, 2, 3, 4, 5 and 8 with Ta at 52°C while reaction 2 uses primers for segment 6, 7, 9 and 10 with Ta at 60°C Then, various PCR conditions were tested by varying the dNTPs (200 nM - 500 nM), MgSO_4_ (1.6 mM - 1.8 mM), enzyme (1 - 2.5 volume) and primer concentrations (100 - 300 nM) to obtain optimal PCR outcomes. Nine RNA templates were used in this assay (Table 2).

### Nanopore sequencing

PCR products were pooled (10 PCR reactions and 2 PCR reactions were pooled for the singleplex and multiplex protocol, respectively) followed by PCR clean up using NucleoSpin Gel and PCR Clean-up column (Macherey-Nagel) and quantification with Qubit dsDNA Broad Range kit (Invitrogen). Approximately 250 ng of the purified and pooled amplicons was used as the template for library preparation using the native barcoding expansion 1-12 kit (EXP-NBD104) according to the manufacturer’s instructions. The prepared library was loaded onto a R9.4.1 Flongle and sequenced for 24 hours. Basecalling of the fast5 raw signals used Guppy v4.4.1 in super accuracy mode to generate the fastq sequences for subsequent bioinformatics analysis.

### Reference-based genome assembly of TiLV samples

Raw reads were quality- and length-filtered using NanoFilt (qscore > 9 and length > 250bp). The raw and filtered read statistics were generated using seqkit v.2.1.0. Reference-based genome assembly of the TiLV was performed according to the ARTIC pipeline (https://github.com/artic-network/fieldbioinformatics) ^15^. This pipeline is an open-source software that integrates a series of tools for base-calling, quality control, read trimming, reference-based mapping, variant calling, consensus sequence generation, and annotation. Briefly, the filtered reads were aligned to the reference TilV genome using Minimap2 v2.17 ^16^ followed by variant calling using Medaka (r941_min_sup_g507) (https://github.com/nanoporetech/medaka). The variants identified were subsequently filtered based on several criteria, including the quality score, depth of coverage, strand bias, and frequency of occurrence. In addition, genomic regions with read depth of lower than 20x were masked prior to generating the final consensus sequence for each sample. Each assembled viral segment from each sample was analyzed with QUAST v5 ^17^ to calculate the percentage of the assembled viral genome that is represented by gaps (Ns), providing insights into the PCR and pooling efficiency.

### Phylogenetic analysis

The assembled viral segments with less than 20% gap were selected and combined with publicly available TiLV genomes for phylogenetic analysis (Table 2). The DNA sequences of the viral genome segments from each sample were extracted and grouped based on their segment number followed by alignment with MAFFT v8 (--adjustdirection --maxiterate 1000 --localpair) ^18^. All 10 individual alignments were subsequently concatenated and used to reconstruct a maximum likelihood tree using FastTree 2 ^19^. The resulting tree was visualized and annotated using FigTree v1.4.4 (http://tree.bio.ed.ac.uk/software/figtree/).

### Code availability

The Linux scripts used to generate raw fastq files, assembled genomes and phylogenetics tree are publicly available in the Zenodo.org dataset (https://zenodo.org/record/7851622).

## Results

### Ten primer pairs for the recovery of complete TiLV genome from various isolation sources

A total of 10 primer pairs were designed with their PCR condition optimized (Table 1 and Supplemental Table 1) to amplify the complete segment of one of the ten TiLV genomic segments. Intact and specific band corresponding to the respective size of the TiLV genomic segments were successfully obtained when the total RNA extracted from TiLV-infected tilapia, TiLV-infected E-11 cell line, and concentrated pond water sample were used as the template for RT-PCR (Figure 1). However, the PCR band intensity for segment 4 (1,250 bp) of the water samples is substantially lower compared to the other segments, requiring another round of PCR (Table 3, Supplemental Table 1).

**Table 3.**
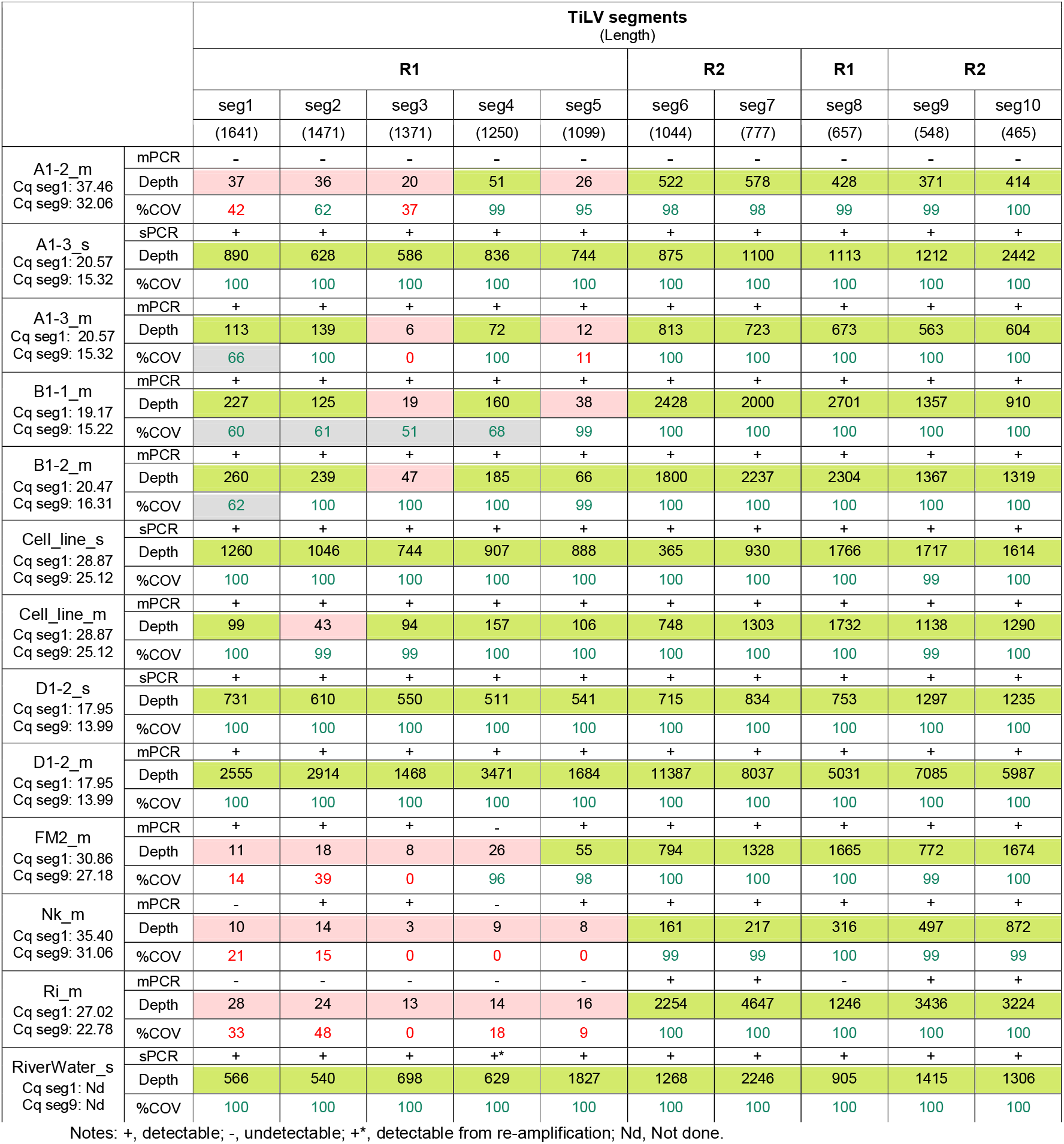
PCR outcome, sequencing, and alignment statistics of 10 individual TiLV segments for each sample used in this study. Viral segments (seg1-5 and seg8) and (seg6, 7, 9, 10) were amplified in two multiplex reactions, reaction 1 (R1) and reaction 2 (R2), respectively. The Cq values from the qPCR detection of segment 1 (Cq seg1) and segment 9 (Cq seg9) were shown below each sample. +/-in the sPCR (singleplex PCR) and mPCR (multiplex PCR) row indicates presence (+) or absence (-) of visible PCR band representing the respective viral segment. %COV (% coverage) indicates the percentage of bases in the assembled contig that consists of a non-ambiguous base; %COV<50 in red and > 50 in green; Depth indicates sequencing depth (or read depth); Depth < 50 are colored in orange and > 50 in green.

**Figure 1.**
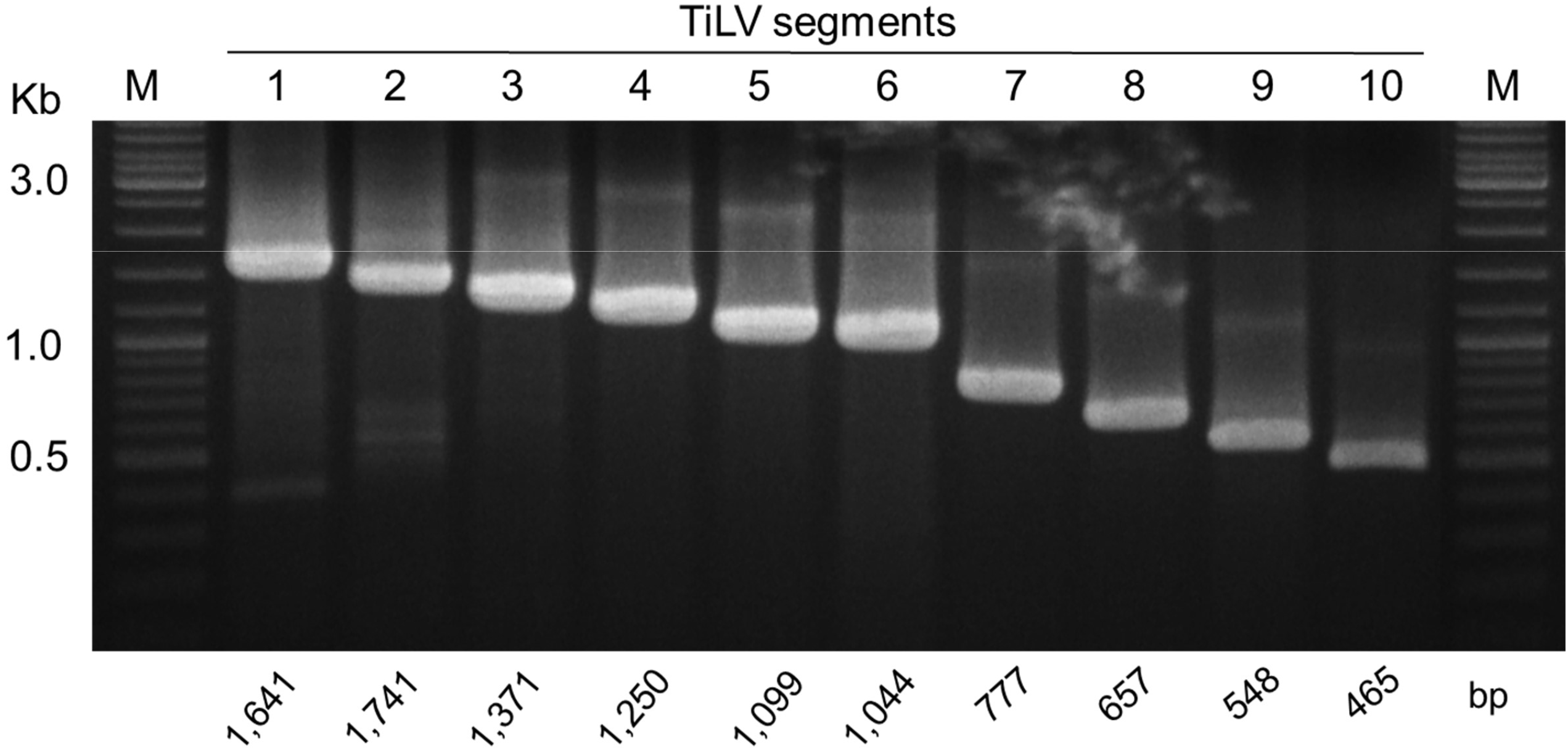
Gel electrophoresis results of one-step RT-PCR amplification of 10 genomic segments of TiLV. Representative results from sample D1-2 are shown. A 1% agarose gel was used to visualize the PCR products, with expected band sizes indicated at the bottom of the gel. M, DNA marker (New England Biolabs).

### A streamlined two-tube multiplex RT-PCR for TiLV genome amplification

To minimize the risk of human error associated with handling multiple singleplex PCR reactions (10 per template), and to reduce chemical costs, a two-tube multiplex RT-PCR was designed (Supplemental Table 1). The addition of MgSO_4_ (increased magnesium ion concentration) was crucial for improved sensitivity (stronger band intensity) while an increase in dNTP concentration does not improve PCR efficiency (Supplemental Figure 1). In addition, increasing the amount of RT/Taq enzyme mix was also shown to slightly improve over band intensity (Supplemental Figure 1). As a result, the concentration of MgSO_4_ and RT/Taq enzyme mix was increased in further multiplex RT-PCR assays (Supplemental Figure 2). After applying the final mPCR conditions to the 10 RNA templates, it was not surprising to observe that samples with high TiLV loads (as determined by qPCR assays) (Table 2) produced the expected six bands and four bands in mPCR reaction 1 and 2, respectively (Figure 2, Table 3). These samples included A1-3, B1-1, B1-2, D1-2, FM2, and the cell line. In contrast, samples NK and Ri, which had lower TiLV loads, showed some missing amplicons, while sample A1-2 exhibited no observable bands in either multiplex RT-PCR reactions (Figure 2, Table 3). It is important to note that mPCR reaction 1 is less sensitive than mPCR reaction 2 (Figure 2) and inconsistently produces observable bands when the Cq value of the tested sample exceeds 19.

**Figure 2.**
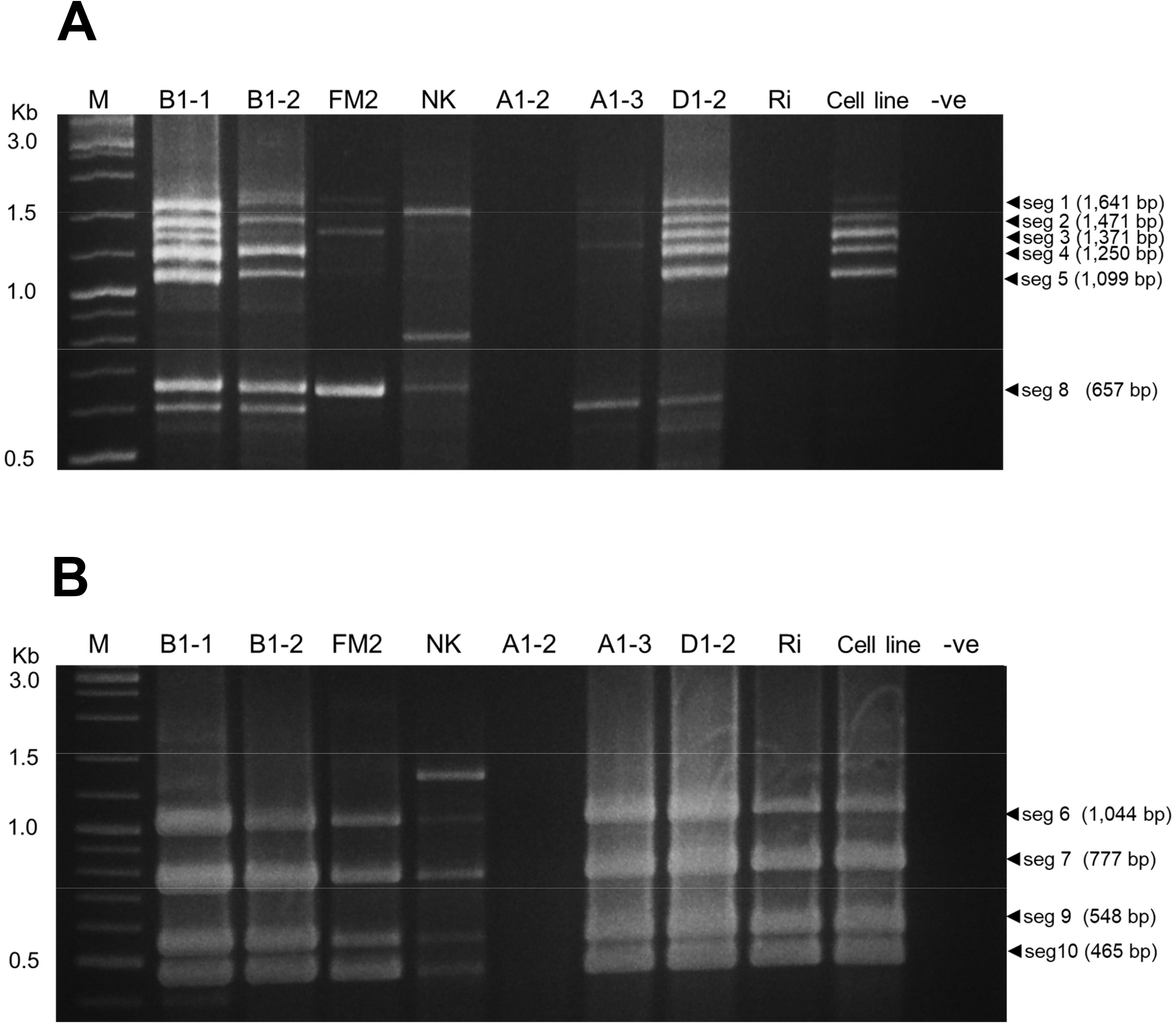
Amplification results of multiplex PCR (mPCR) for TiLV segments. Two separate reactions (Reaction#1 and Reaction#2) were used to amplify 10 TiLV segments. Reaction#1 amplified segments 1, 2, 3, 4, 5, and 8 (Fig 2A), while Reaction#2 amplified segments 6, 7, 9, and 10 (Fig 2B). A 2-log DNA marker (New England Biolabs) was used to visualize the PCR products. -ve, no template control. Codes of samples are listed in Table 2.

### Rapid and on-site sequencing of the TiLV genome using Oxford Nanopore

A total of 413,379 demultiplex raw reads with an accumulative length of 238,983,973 bp were generated from the Flongle sequencing runs (Supplemental Table 2). After filtering (qscore > 9, length > 250 bp), only 194,564 reads (147,415,018 bp) remain. On average more than 35% data reduction was observed across the samples with sample A1-2 showing the largest reduction (67.7%, from 28 Mb to 9.2 Mb) in the amount of usable data. Overall, the Arctic-based reference genome assembly could successfully assemble the viral segments 6,7,8,9,10 for all samples with more than 99-100% completeness except for samples A1-2_m and Nk_m that showed only a slightly lower completeness of 98% for segments 6 and 7 (Table 3). On the contrary, several segments from the first set of multiplex RT-PCR showed reduced completeness (high % of gaps in sequence) particularly for samples with high Cq. The reduced completeness is a direct result of the low read depth (< 20) observed for the viral segments in the respective samples. Generally, any viral segment with a read depth of more than 50x will produce a highly complete assembly that can be used in subsequent analysis (Table 3).

### High phylogenetic diversity among Thai TiLV strains

The total alignment length after the concatenation of 10 individually aligned TiLV viral segments is 10,396 bp. Using a midpoint rooting approach, multiple clades with high SH-like support values were observed in the maximum likelihood tree (Figure 3). Nanopore-sequenced samples from either the pooled singleplex (N= 4 templates) or multiplex amplicons (N= 9 templates) were always placed in the same cluster, consistent with their identical sample origin (Table 2). This observation suggests that accurate genome sequences can be obtained using either singleplex or multiplex amplicon enrichment methods. TiLV strains from Peru, Ecuador, Israel, and India were clustered together and this subclade subsequently formed a sister group with slightly lower support with two earlier TiLV strains from Thailand isolated in 2013 and 2014 to form Clade A (Figure 3). Clade B consisting entirely of Thai TiLV strains from 2015 to 2016 formed a sister group with Clades A. However, a majority of Thai TiLV strains that were reported in 2018 onwards showed yet another distinct clustering as indicated by their phylogenetic placement in Clades C and E with most of the sequences reported in this study belonging to subclade E1. On the other hand, subclade E2 consists of a mixture of Thai and USA TiLV strains. The currently sampled Bangladeshi TiLV strains consist of only single clade despite being isolated 2 years apart (2017 and 2019) while the Vietnamese strains are highly divergent even between themselves (Pairwise nucleotide similarity of only 92%), possibly representing novel strains of TiLV.

**Figure 3.**
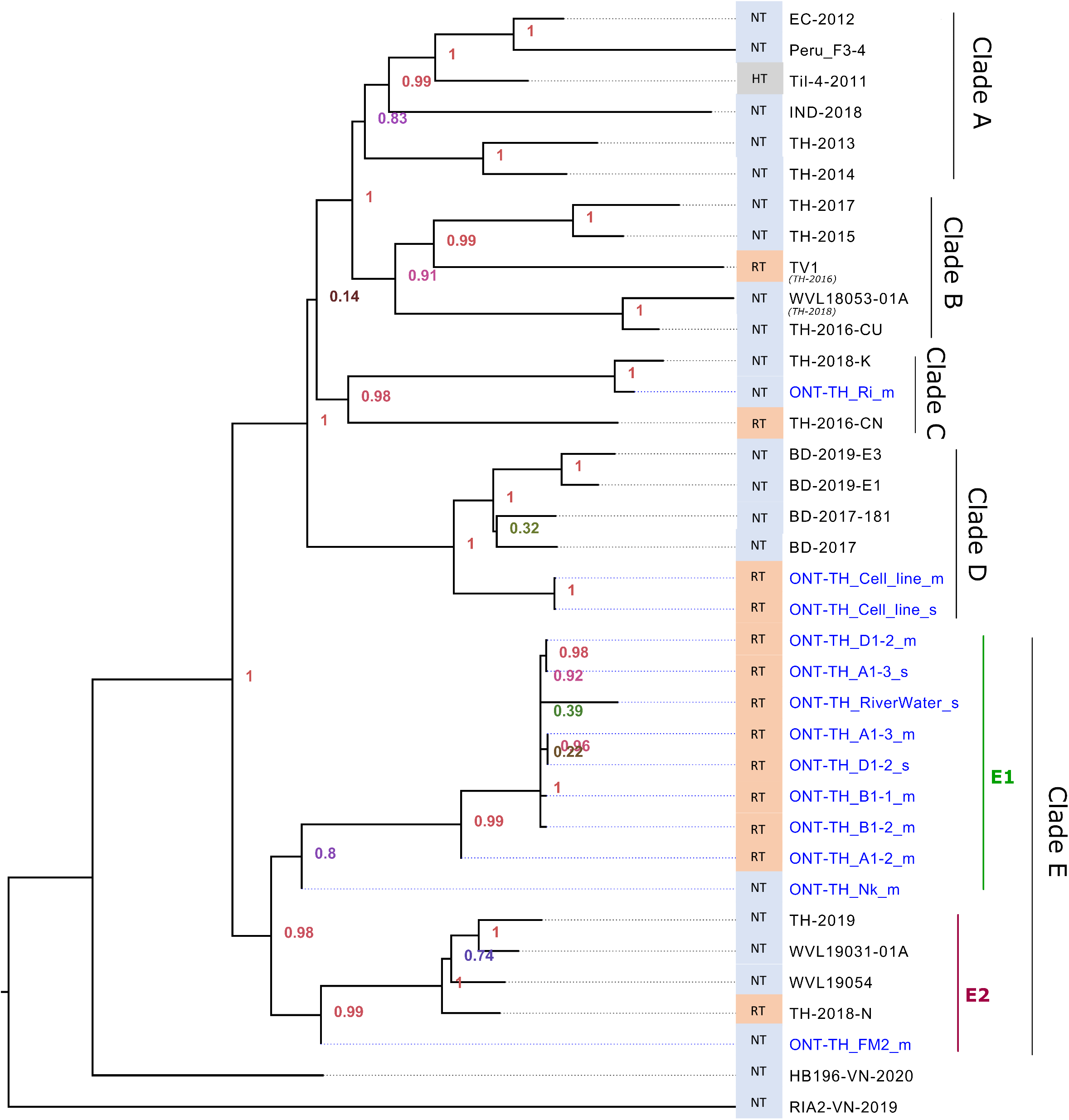
Maximum likelihood tree showing the evolutionary relationships of TiLV strains analyzed in this study. Thirteen samples (10 unique strains) with ONT-TH prefix and publicly available genomes were used. The blue colored tip labels indicate the TiLV strains reported in this study. SH-like local support values and branch length indicate the number of substitutions per site. NT: Nile tilapia; RT: Red tilapia; HT: Hybrid tilapia.

## Discussion

In this study, we report the successful recovery of the complete TiLV genome using a novel approach that combines singleplex PCR and multiplex PCR and Nanopore amplicon sequencing. Our findings indicate that mPCR is particularly effective for samples with high TiLV loads. Therefore, we recommend utilizing mPCR for heavily infected TiLV samples, while sPCR can be employed for lightly infected samples. Our method employs a primer binding region at the terminal 5’ and 3’ ends of each viral segment, which ensures maximal preservation of genetic information. Notably, since the maximum length of the viral segment is less than 1,500 bp, our approach obviates the need for a PCR-tiling strategy typically used for recovering large non-segmented viruses such as ISKNV and SAR-CoV-2 ^12,20^. Moreover, our method offers the added advantage of visualizing PCR efficiency and specificity for each viral segment on a gel, as each fragment has a different size.

Our current multiplex PCR appears to show lower efficiency for Multiplex Set 1, particularly in samples with high Cq values. To improve the multiplex RT-PCR amplification uniformity and efficiency, the Multiplex Set 1 reaction, which amplifies TiLV genomic segments 1-5 and 8, can be further split into two pools (e.g., 1A and 1B) that will amplify an average of three viral segments each. In addition, the use of a more processive High-Fidelity Taq polymerase such as Q5 from New England Biolabs that was currently used for high-degree multiplex tiling PCR of the SAR-CoV19 and ISKNV viral genomes is also worth exploring ^12,15^. It is also worth noting that despite the absence of visible bands for some of the samples, partial or even near-complete genome assembly was still attainable using our sequencing pipeline. It is possible that the amount of PCR product is below the detection limit of gel-staining dye at its loading concentration although it is in fact present in the gel. To streamline future work in high throughput sequencing of TiLV using this approach, gel visualization may be skipped once a lab can consistently reproduce the PCR outcome with evidence from sequencing data.

Nanopore sequencing is an attractive approach for viral amplicon sequencing due to its portability, convenience, and speed ^13^. Our method, which utilizes Nanopore sequencing, eliminates the need for additional fragmentation steps, allowing motor proteins to be directly ligated to amplicons for native sequencing. On the same day, tens of samples can be prepared and sequenced, and the low computing requirements of the ARTIC protocol enable swift genome assembly on a laptop computer, without requiring access to a dedicated server. To further streamline TiLV genome sequencing on the Nanopore platform, we suggest designing multiplex primers that incorporate a partial adapter suitable for Nanopore sequencing ^21^. This enables cost-effective PCR-based barcoding that is both efficient and scalable. In cases of low data output, samples can be re-pooled and sequenced on a separate flow cell to achieve the necessary sequencing depth for genome assembly.

By utilizing R9.4.1 sequencing chemistry with super accuracy mode and implementing the ARTIC pipeline, we successfully recovered TiLV genomes that are highly suitable for phylogenetic inference. Our study revealed the presence of TiLV in both fish and environmental water samples from the same farm, which clustered together in Clade E1. Our approach, combining the previously reported water sample concentration method ^14^ with a multiplex RT-PCR amplicon-based Nanopore sequencing strategy, allowed for direct recovery of TiLV genomes from water samples. This innovative method has significant implications for non-lethal, environmental DNA/RNA monitoring, as it eliminates the need for sacrificing fish for genomic analysis.

Our analysis suggests that, in addition to country of origin, the genetic background of the hosts may also contribute to the clustering patterns observed within Clade E. With few exceptions, our results indicate phylogenetic grouping of Thai TiLV strains (E1: red tilapia, E2: Nile tilapia), suggestive the likelihood of multiple introductions into the country or rapid viral evolution. The presence of the Thai isolates in multiple clusters indicates a significant genetic diversity within the virus. RNA viruses are known for their high mutation rate attributed to the absence of proofreading ability in RNA polymerases ^22^, allowing them to undergo rapid evolutionary changes.

Furthermore, our findings based on the current genomic sampling contradict the initial hypothesis previously put forth on Tilapia trade movement, which was based on a small genome-based phylogenetic tree with limited supported clustering of Bangladeshi and Thai TiLV strains ^9^. Specifically, we found no grouping of Thai strains within the Bangladesh clade (Figure 3, Clade D), thereby reducing support for the previously proposed hypothesis.

Although viral whole genome sequencing of TiLV is now technically feasible, the current representation of its genome in public databases is limited, making it difficult to infer its evolutionary relationships. Given the significant impact of TiLV on the tilapia aquaculture industry, there is a critical need for more robust genomic surveillance to facilitate better management and tracking. Our method can be used in future studies to generate more representative genomes from Vietnam. Our proposed multiplex PCR Nanopore-based amplicon sequencing approach offers a promising solution, as it enables cost-effective and high-throughput sequencing of TiLV virus genomes. This strategy is poised to revolutionize the field of advanced diagnostics and surveillance of multiple pathogens concurrently from biological samples of animals as well as environmental DNA/RNA of pathogens in water, within a single assay. This strategy eliminates the need for separate reactions and reduces the overall cost and time required for sequencing multiple samples. We anticipate that our approach will provide a valuable resource for ongoing efforts to understand the molecular epidemiology and evolution of TiLV, with important implications for disease control and prevention (e.g., vaccination).

## Data Availability

The datasets generated during and/or analyzed during the current study that support our findings are available at the following links: demultiplexed FastQ files for all ten samples can be found under BioProject PRJNA957495 with the corresponding BioSample accession <u>SAMN34257318</u> (B1-1), <u>SAMN34257319</u> (B1-2), <u>SAMN34257320</u> (FM2), <u>SAMN34257321</u> (Nk), <u>SAMN34257322</u> (A1-2), <u>SAMN34257323</u> (A1-3), <u>SAMN34257324</u> (D1-2), <u>SAMN34257325</u> (Ri), <u>SAMN34257326</u> (Cell_line), <u>SAMN34257327</u> (RiverWater).

The intermediary files generated during the bioinformatic analyses are publicly available in the Zenodo.org dataset (https://zenodo.org/record/7851622).

## Acknowledgments

This research was undertaken as part of the CGIAR Initiatives on Aquatic Foods led by WorldFish, and the One CGIAR Initiative “Protecting Human Health Through a One Health Approach”, supported by contributors to the CGIAR Trust Fund: https://www.cgiar.org/funders/. Funding provided support in the form of salary for authors [J.D.D; C.V.M], travel, laboratory consumables and analytical costs. Funders were not involved in the conceptualization, design, data collection, analysis, decision to publish, or preparation of the manuscript.

## Competing interests

The authors declare no competing interests.

**Supplementary Figure 1.**
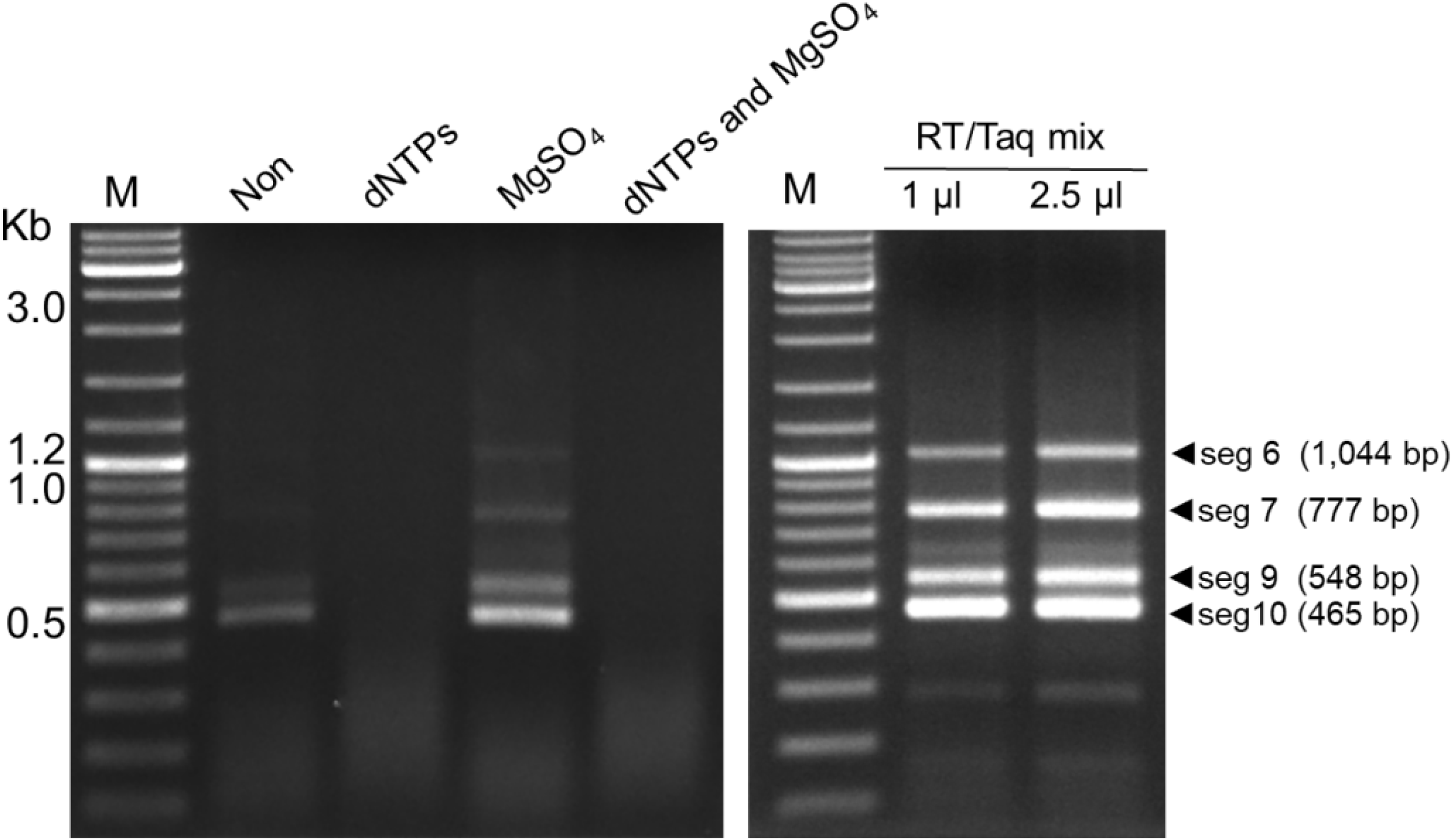
Multiplex RT-PCR condition optimization. The original reaction (Non) and modified reactions with additions of dNTPs, MgSO_4_, dNTPs + MgSO_4_, and RT/Taq enzyme mix were compared. Representative results from mPCR reaction 2 are shown. A DNA marker (New England Biolabs) was used to visualize the PCR products. RNA sample Ri (Table 2) was used in this trial.

**Supplementary Figure 2.**
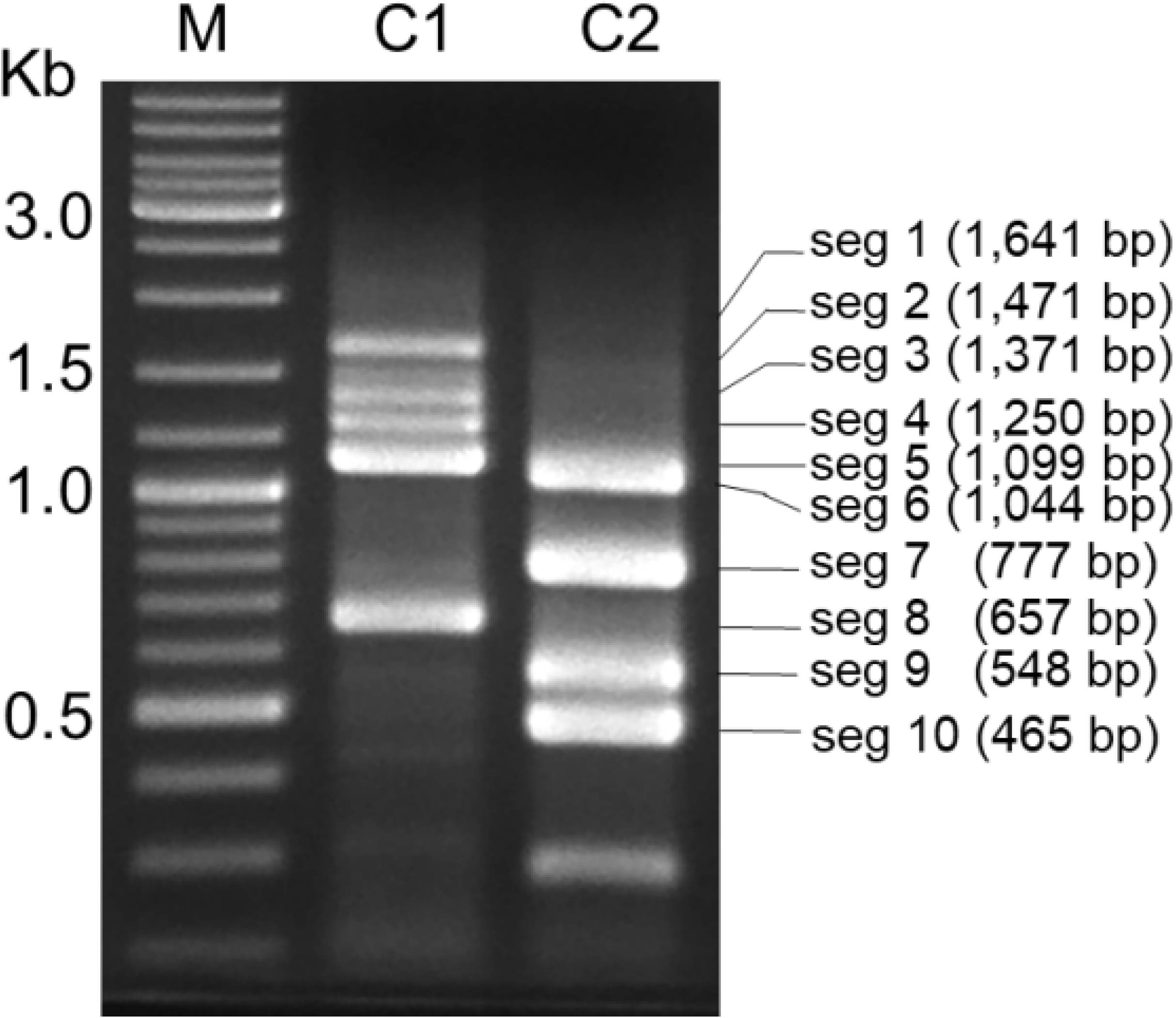
Multiplex RT-PCR amplification of TiLV segments. Two sets of reactions were used to amplify 10 TiLV segments using TiLV RNA templates. Conditions C1 and C2 were used for Reaction 1 and Reaction 2, respectively. C1 amplified segments 1, 2, 3, 4, 5, and 8, while C2 amplified segments 6, 7, 9, and 10. A DNA marker (New England Biolabs) was used to visualize the PCR products. RNA sample Ri (Table 2) was used in this experiment.

## References

1. Aich, N., Paul, A., Choudhury, T. G. & Saha, H. Tilapia Lake Virus (TiLV) disease: Current status of understanding. Aquaculture and Fisheries 7, 7–17 (2022).

2. Jansen, M. D., Dong, H. T. & Mohan, C. V. Tilapia lake virus: a threat to the global tilapia industry? Reviews in Aquaculture 11, 725–739 (2019).

3. Tran, T. H. et al. Tilapia Lake Virus (TiLV) from Vietnam is genetically distantly related to TiLV strains from other countries. Journal of Fish Diseases 45, 1389–1401 (2022).

4. He, T. et al. Identification and pathogenetic study of tilapia lake virus (TiLV) isolated from naturally diseased tilapia. Aquaculture 565, 739166 (2023).

5. Hounmanou, Y. M. G. et al. Tilapia lake virus threatens tilapiines farming and food security: Socio-economic challenges and preventive measures in Sub-Saharan Africa. Aquaculture 493, 123–129 (2018).

6. Bacharach, E. et al. Characterization of a novel orthomyxo-like virus causing mass die-offs of tilapia. mBio 7, e00431–16 (2016).

7. Surachetpong, W. et al. Outbreaks of Tilapia Lake Virus Infection, Thailand, 2015-2016. Emerg Infect Dis 23, 1031–1033 (2017).

8. Thawornwattana, Y. et al. Tilapia lake virus (TiLV): Genomic epidemiology and its early origin. Transboundary and Emerging Diseases 68, 435–444 (2021).

9. Chaput, D. L. et al. The segment matters: probable reassortment of tilapia lake virus (TiLV) complicates phylogenetic analysis and inference of geographical origin of new isolate from Bangladesh. Viruses 12, 258 (2020).

10. Al-Hussinee, L., Subramaniam, K., Ahasan, M. S., Keleher, B. & Waltzek, T. B.Complete Genome Sequence of a Tilapia Lake Virus Isolate Obtained from Nile Tilapia (Oreochromis niloticus). Genome Announcements 6, 10.1128/genomea.00580-18 (2018).

11. Gallagher, M. D. et al. Nanopore sequencing for rapid diagnostics of salmonid RNA viruses. Sci Rep 8, 16307 (2018).

12. Alathari, S. et al. A Multiplexed, Tiled PCR Method for Rapid Whole-Genome Sequencing of Infectious Spleen and Kidney Necrosis Virus (ISKNV) in Tilapia. Viruses 15, 965 (2023).

13. Delamare-Deboutteville, J. et al. Rapid genotyping of tilapia lake virus (TiLV) using Nanopore sequencing. Journal of Fish Diseases 44, 1491–1502 (2021).

14. Taengphu, S. et al. Concentration and quantification of Tilapia tilapinevirus from water using a simple iron flocculation coupled with probe-based RT-qPCR. PeerJ 10, e13157 (2022).

15. Tyson, J. R. et al. Improvements to the ARTIC multiplex PCR method for SARS-CoV-2 genome sequencing using nanopore. 2020.09.04.283077 Preprint at https://doi.org/10.1101/2020.09.04.283077 (2020).

16. Li, H. Minimap2: pairwise alignment for nucleotide sequences. Bioinformatics 34, 3094–3100 (2018).

17. Gurevich, A., Saveliev, V., Vyahhi, N. & Tesler, G. QUAST: quality assessment tool for genome assemblies. Bioinformatics 29, 1072–1075 (2013).

18. Katoh, K. & Standley, D. M. MAFFT Multiple Sequence Alignment Software Version 7: Improvements in Performance and Usability. Molecular Biology and Evolution 30, 772–780 (2013).

19. Price, M. N., Dehal, P. S. & Arkin, A. P. FastTree 2 – Approximately Maximum-Likelihood Trees for Large Alignments. PLOS ONE 5, e9490 (2010).

20. Lambisia, A. W. et al. Optimization of the SARS-CoV-2 ARTIC Network V4 Primers and Whole Genome Sequencing Protocol. Frontiers in Medicine 9, p(2022).

21. Ezpeleta, J. et al. Robust and scalable barcoding for massively parallel long-read sequencing. Sci Rep 12, 7619 (2022).

22. Steinhauer, D. A., Domingo, E. & Holland, J. J. Lack of evidence for proofreading mechanisms associated with an RNA virus polymerase. Gene 122, 281–288 (1992).

23. Eyngor, M. et al. Identification of a novel RNA virus lethal to tilapia. J. Clin. Microbiol. 52, 4137–4146 (2014).

24. Taengphu, S. et al. Genetic diversity of tilapia lake virus genome segment 1 from 2011 to 2019 and a newly validated semi-nested RT-PCR method. Aquaculture 526, 735423 (2020).

25. Subramaniam, K., Ferguson, H. W., Kabuusu, R. & Waltzek, T. B. Genome sequence of tilapia lake virus associated with syncytial hepatitis of tilapia in an Ecuadorian aquaculture facility. Microbiol Resour Announc 8, e00084–19 (2019).

26. Pulido, L. L. H., Mora, C. M., Hung, A. L., Dong, H. T. & Senapin, S. Tilapia lake virus (TiLV) from Peru is genetically close to the Israeli isolates. Aquaculture 510, 61–65 (2019).

27. Debnath, P. P. et al. Two-year surveillance of tilapia lake virus (TiLV) reveals its wide circulation in tilapia farms and hatcheries from multiple districts of Bangladesh. Journal of Fish Diseases 43, 1381–1389 (2020).

