## Supplemental Tables for "A multiplexed RT-PCR Assay for Nanopore Whole Genome Sequencing of Tilapia lake virus (TiLV)"

**Supplemental Table 1.** Reaction mixture and cycling conditions for TiLV quantification and amplification of TiLV segments.

| **Method** | | **Target** | **Reaction** | **Cycling conditions** | **Reference** |
| --- | --- | --- | --- | --- | --- |
| **Single PCR (sPCR)** | **One-step RT-PCR** | Segment  1 to 10 | A 25 µL one step RT-PCR reaction contained the 150 ng template, 200 nM of each primer, 160 nM of MgSO_4_, 1X Reaction Mix, 1.3 µL SuperScript™ III RT/Platinum™ Taq Mix  Catalog number: 12574026 | Reverse transcription 50°C for 10 min, 94°C for 2 min and 35 cycles of 94°C for 30 s, *52°C or *60°C for 30 s and 72°C for 110 s.  Final extension 72°C for 2 min  *52°C for segment 1, 2, 3, 4, 5, 8  *60°C for segment 6, 7, 9, 10 | This study |
|  | **Re-amplification** | Segment 4 | A 50 µL re-amplification PCR reaction contained the 2 µL of the above RT-PCR product, 200 nM of each primer, 1X Terra PCR Direct Buffer and 1.25 U Terra PCR Direct Polymerase Mix  Catalog number: 639270 | 94°C for 2 min, and 40 cycles of 94°C for 30 s, 50°C for 30 s and 72°C for 110 s.  Final extension 72°C for 2 min | This study |
| **Multiplex PCR (mPCR)** | **cDNA synthesis** | Segment  1 to 10 | **I)** A 12 µL cDNA synthesis reaction contained the 2 µg RNA template and 200 nM of each reverse primer.  **II)** A 40 µL reaction contained the 12 µL of reaction **I)**, 500 nM of dNTPs, 3 mM MgSO_4_, 1X ImProm-II™ 5X Reaction Buffer, 320 U Improm-II™ Reverse Transcriptase and nuclease-free water to a final volume of 40 µL  Catalog number: A3802 | **I)** 70°C for 5 min and 4°C for 5 min  **II)** 25°C for 5 min, 42°C for 60 min and 70°C for 15 min. | This study |
|  | **Reaction# 1** | Segment  1, 2, 3, 4, 5 and 8 | A 50 µL PCR reaction contained the 6 µL cDNA template, primers with indicated concentrations below, 1X Terra PCR Direct Buffer and 1.25 U Terra PCR Direct Polymerase Mix  *100 nM of F/R primer for segment 8  *200 nM of F/R primer for segment 1, 3, 4, 5  *300 nM of F/R primer for segment 2 | 94°C for 2 min, and 35 cycles of 94°C for 30 s, 52°C for 30 s and 68°C for 90 s.  Final extension 68°C for 2 min | This study |
|  | **Reaction# 2** | Segment  6, 7, 9 and 10 | A 50 µL PCR reaction contained the 6 µL cDNA template, 200 nM of each primer, 1X Terra PCR Direct Buffer and 1.25 U Terra PCR Direct Polymerase Mix | 94°C for 2 min, and 35 cycles of 94°C for 30 s, 60°C for 30 s and 68°C for 90 s.  Final extension 68°C for 2 min | This study |

| **Method** | **Target** | **Reaction** | **Cycling conditions** | **Reference** |
| --- | --- | --- | --- | --- |
| **RT- qPCR** | Segment 1 | A 20 µL one step RT-qPCR reaction contained the 200 ng RNA template, 450 nM of TiLV-S1-qF (5’-CAA GTC AGT AGA GAT TGA AAG CT-3’), 450 nM of TiLV-S1-qR (5’-CCC ACT TAC ACA ACG AGG AAT-3’), 150 nM of TiLV-S1-qProbe (5’-6-FAM-TTC CCT CCC AGG CAT GTG GGG A-BHQ-1 -3’)  and 1X qScript XLT 1-Step RT-qPCR ToughMix Low Rox (Quanta Bio)  Catalog number: 95134-500 | Reverse transcription 50°C for 10 min, 95°C for 1 min and 40 cycles of 95°C for 10 s and 58°C for 30 s. | This study |
|  | Segment 9 | A 20 µL one step RT-qPCR reaction contained the 150 ng RNA template, 450 nM of TiLV-S9-qF (5′-CTA GAC AAT GTT TTC GAT CCA G-3′), 450 nM of TiLV-S9-qR (5′-TTC TGT GTC AGT AAT CTT GAC AG-3′), 150 nM of TiLV-S9-qProbe (5′-6-FAM-TGC CGC CGC AGC ACA AGC TCC A-BHQ-1-3′) and 1X qScript XLT 1-Step RT-qPCR ToughMix Low Rox (Quanta Bio)  Catalog number: 95134-500 | Reverse transcription 50°C for 10 min, 95°C for 1 min and 40 cycles of 95°C for 10 s and 58°C for 30 s. | Taengphu et al, 2022 |

**Supplemental Table 2**. Sequencing trimming statistics

| **Samples** | **#_seqs** | **%seq**  **trimmed** | **Sum base-pair (bp)** |  | **%base**  **trimmed** | **min**  **length** | **avg**  **length** | **max**  **lenght** | **N50** |
| --- | --- | --- | --- | --- | --- | --- | --- | --- | --- |
| A1-2_m | 121135 | 82.71% | 28,671,617 |  | 67.74% | 9 | 236.7 | 282415 | 249 |
| A1-2_m | 20948 |  | 9,248,348 |  |  | 250 | 441.5 | 282415 | 420 |
| A1-3_m | 7446 | 41.02% | 4,689,273 |  | 32.37% | 44 | 629.8 | 8173 | 764 |
| A1-3_m | 4392 |  | 3,171,523 |  |  | 250 | 722.1 | 8173 | 769 |
| A1-3_s | 24149 | 49.01% | 18,753,392 |  | 45.18% | 1 | 776.6 | 12945 | 1041 |
| A1-3_s | 12314 |  | 10,281,058 |  |  | 250 | 834.9 | 12945 | 1054 |
| B1-1_m | 20170 | 41.66% | 12,146,167 |  | 31.02% | 41 | 602.2 | 3077 | 749 |
| B1-1_m | 11767 |  | 8,378,540 |  |  | 250 | 712 | 3077 | 759 |
| B1-2_m | 18586 | 40.65% | 12,085,327 |  | 32.54% | 40 | 650.2 | 4132 | 762 |
| B1-2_m | 11030 |  | 8,152,745 |  |  | 250 | 739.1 | 4132 | 766 |
| Cell_line_m | 9923 | 32.40% | 7,192,352 |  | 28.21% | 35 | 724.8 | 4403 | 764 |
| Cell_line_m | 6708 |  | 5,163,134 |  |  | 250 | 769.7 | 4403 | 764 |
| Cell_line_s | 23135 | 43.38% | 18,432,299 |  | 37.87% | 1 | 796.7 | 4235 | 1139 |
| Cell_line_s | 13098 |  | 11,452,457 |  |  | 250 | 874.4 | 4034 | 1164 |
| D1-2_m | 82519 | 32.29% | 66,056,539 |  | 28.03% | 27 | 800.5 | 5631 | 1030 |
| D1-2_m | 55870 |  | 47,541,378 |  |  | 250 | 850.9 | 5631 | 1032 |
| D1-2_s | 18408 | 50.21% | 14,741,036 |  | 46.48% | 1 | 800.8 | 15888 | 1066 |
| D1-2_s | 9165 |  | 7,889,536 |  |  | 250 | 860.8 | 15888 | 1072 |
| FM2_m | 11228 | 36.86% | 7,137,381 |  | 31.05% | 41 | 635.7 | 3989 | 745 |
| FM2_m | 7089 |  | 4,921,552 |  |  | 250 | 694.3 | 3989 | 749 |
| Nk_m | 25098 | 47.66% | 12,486,929 |  | 34.03% | 33 | 497.5 | 4476 | 683 |
| Nk_m | 13136 |  | 8,237,670 |  |  | 250 | 627.1 | 4433 | 741 |
| Ri_m | 27610 | 42.81% | 17,425,255 |  | 34.02% | 1 | 631.1 | 3619 | 767 |
| Ri_m | 15790 |  | 11,496,335 |  |  | 250 | 728.1 | 3337 | 769 |
| RiverWater_s | 23972 | 44.70% | 19,166,406 |  | 40.10% | 29 | 799.5 | 14266 | 1040 |
| RiverWater_s | 13257 |  | 11,480,742 |  |  | 250 | 866 | 14266 | 1054 |

Highlighted in red, demultiplex raw reads; highlighted in green, after filtering (Qscore > 9, min read length 250 bp; #_seqs, number of sequences
